# Low blood concentration of alcohol enhances activity related to stopping failure in the right inferior frontal cortex

**DOI:** 10.1101/2023.12.10.571028

**Authors:** Jun Shinozaki, Hiroshi Matsumoto, Hidekazu Saito, Takashi Murahara, Hiroshi Nagahama, Yuuki Sakurai, Takashi Nagamine

## Abstract

This study investigated the effects of low doses of alcohol, which are acceptable for driving a car, on inhibitory control and neural processing using the stop-signal task (SST) in 17 healthy right-handed social drinkers. The study employed simultaneous functional magnetic resonance imaging and electromyography (EMG) recordings to assess behavioural and neural responses under conditions of low-dose alcohol (breath-alcohol concentration of 0.15 mg/L) and placebo. The results demonstrated that even a small amount of alcohol consumption prolonged Go reaction times in the SST and modified stopping behaviour, as evidenced by a decrease in the frequency and magnitude of partial response EMG that did not result in button pressing during successful inhibitory control. Furthermore, alcohol intake enhanced neural activity during failed inhibitory responses in the right inferior frontal cortex, suggesting its potential role in behavioural adaptation following stop-signal failure. These findings suggest that even low levels of alcohol consumption within legal driving limits can greatly impact both the cognitive performance and brain activity involved in inhibiting responses. This research provides important evidence on the neurobehavioural effects of low-dose alcohol consumption, with implications for understanding the biological basis of impaired motor control and decision-making and potentially informing legal guidelines on alcohol consumption.

## INTRODUCTION

Alcohol is a drug that can alter various human functions. Several studies have shown that both high alcohol concentrations and low legally permissible concentrations can cause functional changes. For example, decreases in vigilance and spatial information processing have been reported at blood alcohol concentrations (BACs) as low as 0.03% (an estimated breath-alcohol concentration [BrAC] of 0.15 mg/L) (Koelega 1995). At a BAC of 0.05%, simple and choice reaction times (RTs) were prolonged (Tzambazis and Stough 2000), while at a BAC of 0.0054%, working memory function was impaired (Scheel et al. 2013). A relatively low BAC of 0.045% suppressed the P3a response to novel auditory stimuli (Marinkovic et al. 2001). These findings imply that alcohol consumption in low quantities may have a greater impact on various human functions than previously thought.

Inhibitory control is a cognitive process that prevents inappropriate and undesirable behaviours. This ability is essential for controlling impulsivity and impulse responses and choosing appropriate actions in response to a situation; it allows us to stop our behaviour in the face of sudden danger. The right inferior frontal cortex (rIFC) is a critical brain region involved in the inhibitory control of movement (see Aron et al. 2014 for a review). Damage to the rIFC affects stopping behaviour (Aron et al. 2003; Clark et al. 2007), and transcranial magnetic stimulation (TMS) studies have shown that it delays stopping in the human stop-signal task (SST) (Mulvihill, Skilling, and Vogel-Sprott 1997; de Wit, Crean, and Richards 2000; Chambers et al. 2006, 2007; Loeber and Duka 2009; Verbruggen et al. 2010). According to a meta-analysis of neuroimaging studies, the rIFC is a key region involved in response inhibition (Levy et al. 2011).

The SST is a behavioural measure of response inhibition (Logan and Cowan 1984; Verbruggen and Logan 2008). It consists of Go and Stop trials. In the Go trials, a right or left arrow, referred to as the Go signal, is presented visually, and participants are required to respond by pressing a button indicating the direction of the arrow. Occasionally, after the Go signal, an image or sound called the Stop signal is presented and the participants are instructed to withhold their button press response. These are referred to as Stop trials. Stop trials are further subdivided into stop failure and stop success trials based on the participant’s button press response. Even when participants successfully withhold their button press response, a small muscle contraction may occur, referred to as partial response electromyography (prEMG). PrEMG occurs when muscle activity initiated by the Go signal is rapidly reduced by the Stop signal before the button press response. A review has shown that the frequency of prEMG is inversely correlated with RT in Go trials (Raud et al. 2022). Hence, it is anticipated that the behavioural-EMG link is influenced by the trade-off approach between Go and Stop, as the frequency of prEMG increases when participants try to react rapidly to the Go signal.

Functional magnetic resonance imaging (fMRI) studies using the SST have shown that alcohol consumption prolongs stop-signal reaction time (SSRT), an index of inhibitory control, and decreases neural activity in the rIFC (Gan et al. 2014). However, Gan et al. administered relatively high doses of alcohol compared with those allowed under motor vehicle driving laws and regulations; therefore, it remains unclear whether low-dose alcohol intake, as low as the acceptable limit, affects stop behaviour and stop-related neural activity.

This study aimed to investigate whether low-dose alcohol consumption affects the trade-off strategy between going and stopping and to elucidate the relationship between these strategies and neural activity. Therefore, we performed simultaneous fMRI and EMG measurements during the SST. We adopted a low-dose alcohol intake, or BrAC of 0.15 mg/L, as the minimum legal limit for driving in Japan. Our study aimed to clarify whether low-dose alcohol intake affects stopping behaviour and stopping-related neural activity. Both outcomes provide evidence for the physiological validity of the current law.

## METHODS

### Participants

Twenty-one healthy, right-handed social drinkers (16 men, 5 women) participated, with a mean age of 23 years (range 20-27) and mean weight of 60 kg (range 40-79). There was no history of neurological or psychiatric disorders. All participants abstained from alcohol for 24 hours before the experiment and passed the Kurihama Alcoholism Screening Test (Revised Version: KAST-M for men, KAST-F for women). Written informed consent was obtained from all patients, and the study was approved by the Ethics Committee of Sapporo Medical University.

### Alcohol administration

The first criterion for participating in the study was a BrAC of 0.00 mg/L on arrival at the experiment room (measured using a Dräger Alcotest 3500). Participants who met this requirement participated in the study, which was conducted in a single-blind manner over two separate days, one with alcohol intake and one without alcohol intake (n = 13 with alcohol first, n = 8 with placebo first). Participants consumed 0.3 g/kg-body weight alcohol-mixed or placebo tea within 30 minutes, and BrAC was measured every 5 minutes. fMRI was initiated when the BrAC dropped to no more than 0.20 mg/L. The second criterion for data acceptance was a BrAC of at least 0.10 mg/L at the end of the fMRI session, with a mean estimated BrAC of 0.15 mg/L. Two participants had a BrAC below 0.10 mg/L after the imaging and were not included in further analysis, while two others dropped out, resulting in data from 17 participants being used in the analysis (n = 12 with alcohol first, n=5 with placebo first). The BrAC time courses are shown in the Supplementary Information (Fig. S1).

### Stop-signal task

In each trial, a green arrow pointing to the left or right was displayed in the centre of the screen for 900 ms as the Go cue, and the participants were asked to press the corresponding left or right button using their left or right thumb as quickly as possible. The direction of the arrow was randomised. In 25% of the trials (Stop trials), the Go cue was immediately followed by a red square presented at the screen centre for 900 ms as the Stop cue. The percentage of Stop trials was adopted from previous studies (Rubia et al. 2003; Aron and Poldrack 2006; Li et al. 2006; Leung and Cai 2007; Badry et al. 2009; Cai and Leung 2009; Tabu et al. 2011). Participants were asked to inhibit their button presses during the Stop trials. The onset of the Go cue was jittered from 3 to 5 s, with a fixation cross displayed in the centre throughout the inter-trial interval.

The delay between the onset of the Go and Stop cues was defined as the stop-signal delay (SSD). The SSD was adjusted based on the participant’s performance in the previous Stop trial, with an increase of 50 ms after successful inhibition (Stop-success [SS]) and a decrease of 50 ms after failure (Stop-failure [SF], i.e. pressing the button,). This step-up and step-down staircase ensured convergence to a 50% failure rate by the end of each session, starting at 150 ms from the beginning of each session. RT was defined as the time from the onset of the Go cue to the button press. After each session, the median RT and average SSD (e.g. when the SSDs changed as follows: 150, 100, 150, 200, 150, 200, 150, 100, 150, and 100 ms, the average SSD was 145 ms) were presented to the participants. One session consisted of 30 Stop trials and 90 Go trials, and three sessions were conducted, each of which took approximately 30 minutes.

### EMG recording

Electromyography (EMG) was recorded using the BrainAmp ExG MR system (Brain Products GmbH, Munich, Germany). Two pairs of sintered silver/silver chloride electrodes were bilaterally placed over the thenar muscles. A ground electrode was placed on the surface of either the right or the left elbow. EMG data were collected using Brain Vision Recorder software. EMG signals were transmitted via an optical cable to a PC monitor outside the scanning room. They were filtered with a low cut-off of DC and high cut-off of 1000 Hz and then converted to digital signals with a sampling rate of 5 kHz. The digitisation clock was synchronised with the MRI scanner clock.

For offline analysis, we corrected the EMG signals by excluding scanner artefacts using Brain Vision Analyzer software. The corrected EMG data were then high-pass filtered at 5 Hz (slope: 48 dB/oct), low-pass filtered at 100 Hz (slope: 48 dB/oct), notch filtered at 50 Hz, and downsampled to 500 Hz, and these data were rectified.

### Behavioural analysis

The SSRT was calculated by subtracting the average SSD from the median RT of the correct Go trials, as per Logan’s race model (Logan et al. 1984) (Fig. 1). Incorrect responses in the Go trials were considered error trials, and error rates were calculated. The SS rate was defined as the number of SS trials divided by the total number of Stop trials. A repeated measures ANOVA (rmANOVA) was performed on these behavioural measures, with the within-subject factors ALC (alcohol or placebo) and LtRt (left or right arrow trials). In the case of a significant interaction, behavioural measures were calculated separately for the left and right trials. Otherwise, they were calculated by pooling the left and right trials because the effect of alcohol should be similar across these trials. The effects of alcohol on these parameters were determined by two-tailed paired t-tests.

**Figure 1:**
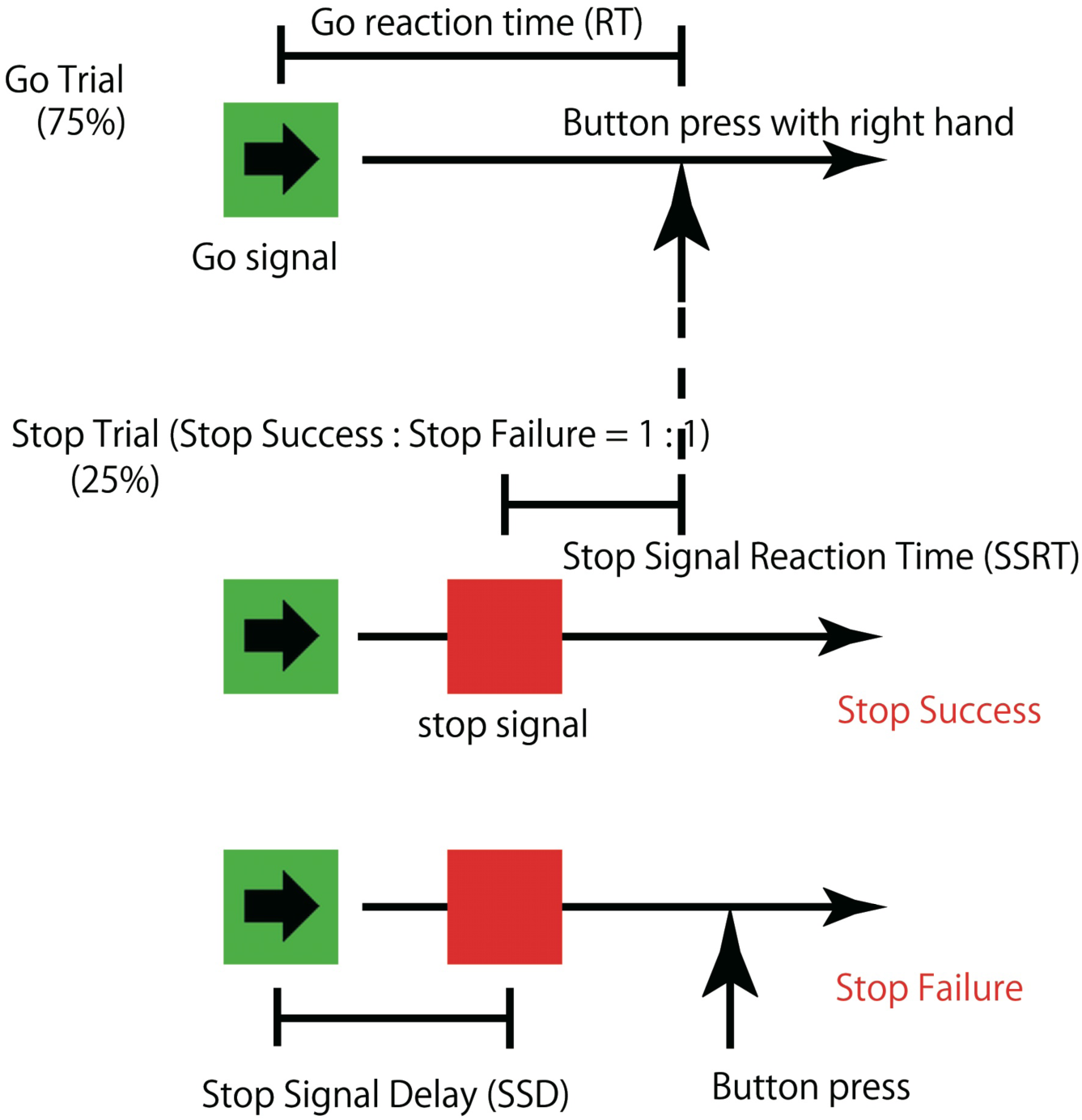
During the stop-signal task (SST), a green arrow appeared for 900 ms and participants pressed the corresponding button quickly. In 25% of the trials, a red square followed as a Stop signal. The timing of the Stop signals varied. Sessions included 30 Stop and 90 Go trials, lasting 30 minutes. The stop-signal delay (SSD) was adjusted by 50 ms after each trial to achieve a 50% inhibition rate. Reaction times were measured, and average SSD and stop-signal reaction time (SSRT) were calculated post-session.

Additionally, we evaluated the RT for SS/SF. It is known that Go RT increases after SF trials (Rabbitt 1966; Zhang et al. 2017). Thus, we compared Go RTs just before and after a Stop signal by subtracting the former from the latter, trial-by-trial, in each participants to determine whether it was altered by the failure or success of the Stop signals. The median of the subtracted data among the SF trials was called post-error slowing (PES) (Hajcak et al. 2003; Zhang et al. 2017). Similarly, this was referred to as post-success slowing (PSS) in SS trials (Zhang et al. 2017). A one-sample t-test was performed on PES and PSS and compared to zero under both alcohol and placebo conditions to evaluate whether slowing occurred. Additionally, rmANOVA was performed on PES and PSS with the within-subject factors SF/SS and ALC.

The EMG data were analysed in two ways. First, partial response EMG (prEMG) (Raud et al. 2022) changes that occurred even in successful response inhibition (SS trials) were evaluated. Significant prEMG was defined as a maximal power between 100 ms and 500 ms after Go cue onset that exceeded 10 times the standard deviation of power in the 300 ms period before onset. PrEMG frequency was calculated as the number of trials with a significant prEMG divided by the number of SS trials. A two-tailed paired t-test was performed on prEMG frequency to evaluate the effects of alcohol consumption. Second, EMG power was analysed using the area under the curve (AUC). To account for individual and electrode position variations, EMG power was defined as the AUC value between 0 and 1000 ms after Go cue onset, divided by the baseline period of 300 ms before onset in a trial-by-trial manner. The mean EMG power throughout the scan was calculated for each category (left Go, left SS, left SF, right Go, right SS, right SF in alcohol and placebo conditions) for each subject, excluding the Go-wrong trials. To account for individual EMG differences, SS and SF EMG powers were divided by the Go EMG power individually, referred to as “normalized EMG magnitude”. The rmANOVA was performed on these normalised EMG magnitudes using the within-subject factors of LtRt, SF/SS, and ALC.

### MRI acquisition

The experiments were conducted using a 3-Tesla scanner equipped with an 8-ch phased array coil (GE Healthcare, Signa). Functional images were acquired using a T2*-weighted gradient-echo echo-planar imaging sequence. The image acquisition parameters were as follows: repetition time (TR) = 2000 ms, echo time (TE) = 20 ms (Shinozaki et al. 2013), flip angle = 90°, field of view (FOV) = 192 mm, matrix = 64 × 64, and 38 ascending slices of 3 mm thickness with a 0.6 mm gap. T1-weighted images were acquired as follows: TR = 5.764 ms, TE = 2.016 ms, inversion time = 400 ms, FOV = 256 mm, matrix = 256 × 256, 1-mm thickness slices (1 mm cubic voxels).

### fMRI data analysis

The imaging data were analysed with SPM8 using a general linear model (Friston et al. 1994). These images were spatially realigned to the mean image to correct for head movements, corrected for slice timing, and co-registered with the individual T1 images. The co-registered T1 image was spatially normalised to a Montreal Neurological Institute (MNI) T1 template (Evans et al. 2005), and the normalised parameters were then applied for the normalisation of the functional images. The normalised functional images were smoothed with an isotropic Gaussian kernel of 6 mm full-width at half-maximum (FWHM). Each of the six categories (left Go, left SF, left SS, right Go, right SF, and right SS) and six head-movement parameters was modelled separately as a regressor for the first-level analysis.

Regarding Go-wrong (error) trials in the Go trials, the left and right Go-wrong regressors were added as parametric modulations for each Go regressor. This analysis was performed for each subject to test the correlation between the MRI signals and the delta function convolved with the canonical haemodynamic response function. Global signal normalisation was not performed during the sessions. Low-frequency noise was removed using a high-pass filter with a cut-off of 128 s, and serial correlation was adjusted using an AR(1) model. Parameter estimates reflecting the magnitude of the correlation between the signals and model of interest were calculated. These were then used for the subsequent second-level random-effects model analysis. Group-level statistical parametric maps were created using a 2 × 2 × 3 factorial design with the within-subject factors ALC (alcohol condition, placebo condition), LtRt (left arrow trials, right arrow trials), and Go/SF/SS (Go, SF, SS). These results are shown at a height threshold of p < 0.001 (uncorrected) with cluster-level p < 0.05, family-wise error (FWE) corrected for the whole brain, and overlaid on the MNI template brain.

## RESULTS

### Behavioural results

As mentioned in the Methods section, to evaluate whether the behavioural measures could be divided into right and left arrow trials, the interaction between ALC and LtRt was investigated using rmANOVA. The rmANOVA (ALC × LtRt) showed a significant interaction on normalized EMG magnitude (F_1, 15_ = 5.869, p = 0.029), while it did not show any significant interactions on the other metrics (F_1, 16_ = 4.319, p = 0.054 for RT; F_1, 16_ = 0.002, p = 0.964 for Go error rate; F_1, 16_ = 0.327, p = 0.575 for SSD; F_1, 16_ = 0.150, p = 0.703 for SSRT; F_1, 16_ = 0.889, p = 0.360 for SS rate). The prEMG frequency did not show a significant interaction (F_1, 16_ = 1.948, p = 0.183). Therefore, we reported the normalised EMG magnitudes in the left and right arrow trials separately and pooled them for other behavioural measures.

RTs were significantly longer in the alcohol condition than in the placebo condition (t_16_ = 2.530, d = 0.492, p = 0.022). The error rate in the Go trials under both the alcohol and placebo conditions was low (approximately 1%), and there was no significant difference between the two conditions (t_16_ = 0.774, d = 0.185, p = 0.450). No significant difference in SSRT was observed between the alcohol and placebo conditions (t_16_ = 0.369, d = 0.051, p = 0.717), nor in the average SSD (t_16_ = 2.018, d = 0.409, p = 0.061). The SS rate in the Stop trials did not show a significant difference from the ideal value of 50% (p > 0.098 in all subjects in the chi-square test), and there was no significant effect of alcohol (t_16_ = 1.410, d = 0.367, p = 0.178) (Table 1).

**Table 1.** Behavioural results. *, effect of alcohol is significant at p < 0.05 (two-tailed paired t-test). S.E.M., standard error off the mean; RT, reaction time; SSRT, stop-signal reaction time; SSD, stop-signal delay.

| Parameters | Condition (mean ± S.E.M.) | | |
| --- | --- | --- | --- |
|  | Alcohol | Placebo | p-value |
| RT (ms) | 415.6 ± 8.2* | 396.9 ± 10.2 | 0.022 |
| Go error rate | 0.014 ± 0.004 | 0.011 ± 0.003 | 0.450 |
| SSRT (ms) | 238.1 ± 6.3 | 236.8 ± 6.4 | 0.717 |
| Average SSD (ms) | 177.5 ± 9.2 | 160.1 ± 11.4 | 0.061 |
| Stop-success rate | 0.53 ± 0.01 | 0.51 ± 0.01 | 0.178 |
| Stop-fail RT (ms) | 386.8 ± 6.6 | 377.1 ± 8.2 | 0.237 |

To investigate whether Go RTs increased after SF and SS, we conducted a one-sample t-test with a Bonferroni correction (corrected p = 0.0125). PES was significantly greater than zero only in the alcohol condition (mean = 17.4 ms, t_16_ = 4.270, p = 0.001), while PES in the placebo condition and PSS in the alcohol and placebo conditions did not show significant difference from zero (mean = 10.6 ms, t_16_ = 2.378, p = 0.015 for PES in placebo condition; mean = 8.9 ms, t_16_ = 2.332, p = 0.017 for PSS in alcohol condition; mean = 4.5 ms, t_16_ = 1.295, p = 0.107 for PSS in placebo condition) (Table 2). To reveal the effect of alcohol on slowing, an rmANOVA was conducted on the PES and PSS. There was no significant interaction between SF/SS and ALC (F_1, 16_ = 0.132, p = 0.721), whereas the main effect of ALC was significant (F_1, 16_ = 5.336, p = 0.035; slowing was greater in the alcohol condition than in the placebo condition). The main effect of SF/SS was not significant (F_1, 16_ = 2.356, p = 0.144).

**Table 2.** Post-error slowing (PES) and post-success slowing (PSS). *, effects of slowing are significant at p < 0.0125 (one-tailed one-sample t-test [Bonferroni-corrected]). cond., condition; S.E.M., standard error of the mean.

| Parameters | Slowing (mean ± S.E.M.) | p-value |
| --- | --- | --- |
| **Post-Error Slowing** |  |  |
| PES of alcohol cond. (ms) | 17.4 ± 4.1* | 0.001 |
| PES of placebo cond. (ms) | 10.6 ± 4.4 | 0.015 |
| **Post-Success Slowing** |  |  |
| PSS of alcohol cond. (ms) | 8.9 ± 3.8 | 0.017 |
| PSS of placebo cond. (ms) | 4.5 ± 3.5 | 0.107 |

EMG leakage in the SS trials was obtained during fMRI from 16 participants except one participant, whose data were missing because of machine trouble. prEMG frequency was 0.10 under the alcohol condition, which was significantly lower than 0.19 in the placebo condition (t_15_ = 2.296, p = 0.036). The rmANOVA of the normalised EMG magnitude showed a significant main effect of ALC (F_1, 15_ = 6.025, p = 0.027; EMG under the alcohol condition was lower than that under the placebo condition). The LtRt × SF/SS × ALC interaction was significant (F_1, 15_ = 13.281, p = 0.002). In the SS right arrow trials, the normalised EMG magnitude was significantly smaller in the alcohol condition than in the placebo condition (F_1, 60_ = 25.471, p < 0.001). In addition, there was no significant difference between the alcohol and placebo conditions in the SS left, SF right, and SF left arrow trials (F_1, 60_ = 0.200, p = 0.656; F_1, 60_ = 0.438, p = 0.511; F_1, 60_ = 0.215, p = 0.644, respectively) (Table 3). Taken together, both the prEMG frequency and magnitude were lower in the SS trials in the alcohol condition than in the placebo condition.

**Table 3.** prEMG frequency and normalized EMG magnitude. Each EMG magnitude was divided by the EMG magnitude of Go trials. *, simple effect of alcohol is significant at p < 0.05 (repeated measures ANOVA). prEMG, partial response electromyography; EMG, electromyography; S.E.M., standard error of the mean.

| Parameters | Condition (mean ± S.E.M.) | | |
| --- | --- | --- | --- |
|  | Alcohol | Placebo | p-value |
| prEMG frequency | 0.10 ± 0.03* | 0.19 ± 0.05 | 0.036 |
| **Normalized EMG magnitude (Left arrow trial)** |  |  |  |
| Stop-success/Go | 0.279 ± 0.030 | 0.291 ± 0.021 | 0.656 |
| Stop-failure/Go | 0.911 ± 0.019 | 0.924 ± 0.013 | 0.644 |
| **Normalized EMG magnitude (Right arrow trial)** |  |  |  |
| Stop-success/Go | 0.184 ± 0.024* | 0.323 ± 0.037 | < 0.001 |
| Stop-failure/Go | 0.914 ± 0.018 | 0.932 ± 0.017 | 0.511 |

### fMRI results

We first investigated whether the effect of LtRt on brain activity was significant. We focused on brain activity related to response inhibition, that is, the interruption of an already initiated Go process (Aron and Poldrack 2006). Thus, we compared the brain activity in the left arrow trials with that in the right arrow trials, in contrast to the SS vs Go. The subtraction between left and right (Lt(SS − Go) − Rt(SS − Go) and vice versa) revealed significant effects of LtRt only in the somatomotor area and cerebellum, and no significant effect occurred in the rIFC (cluster level FWE-corrected p < 0.05). Regarding the contrast of SF vs Go, the subtraction between left and right (Lt(SF − Go) − Rt(SF − Go) and vice versa) did not provide any effects of LtRt in the whole brain (cluster level FWE-corrected p < 0.05). Thus, we pooled the left and right arrow trials for subsequent analysis.

In the comparison of SS and Go (pooling the alcohol/placebo conditions), we observed significant activity in the bilateral parietal cortices, bilateral inferior and superior frontal gyri, insulae, and medial frontal gyri (Fig. 2A, Table 4). To investigate the neural activity related to the failure of response inhibition, we compared SF and Go (pooled alcohol/placebo conditions). This contrast of (SF vs Go) revealed significant activity in the bilateral parietal cortices, inferior and superior frontal gyri, insulae, and anterior cingulate cortices (Fig. 2B, Table 5).

**Figure 2:**
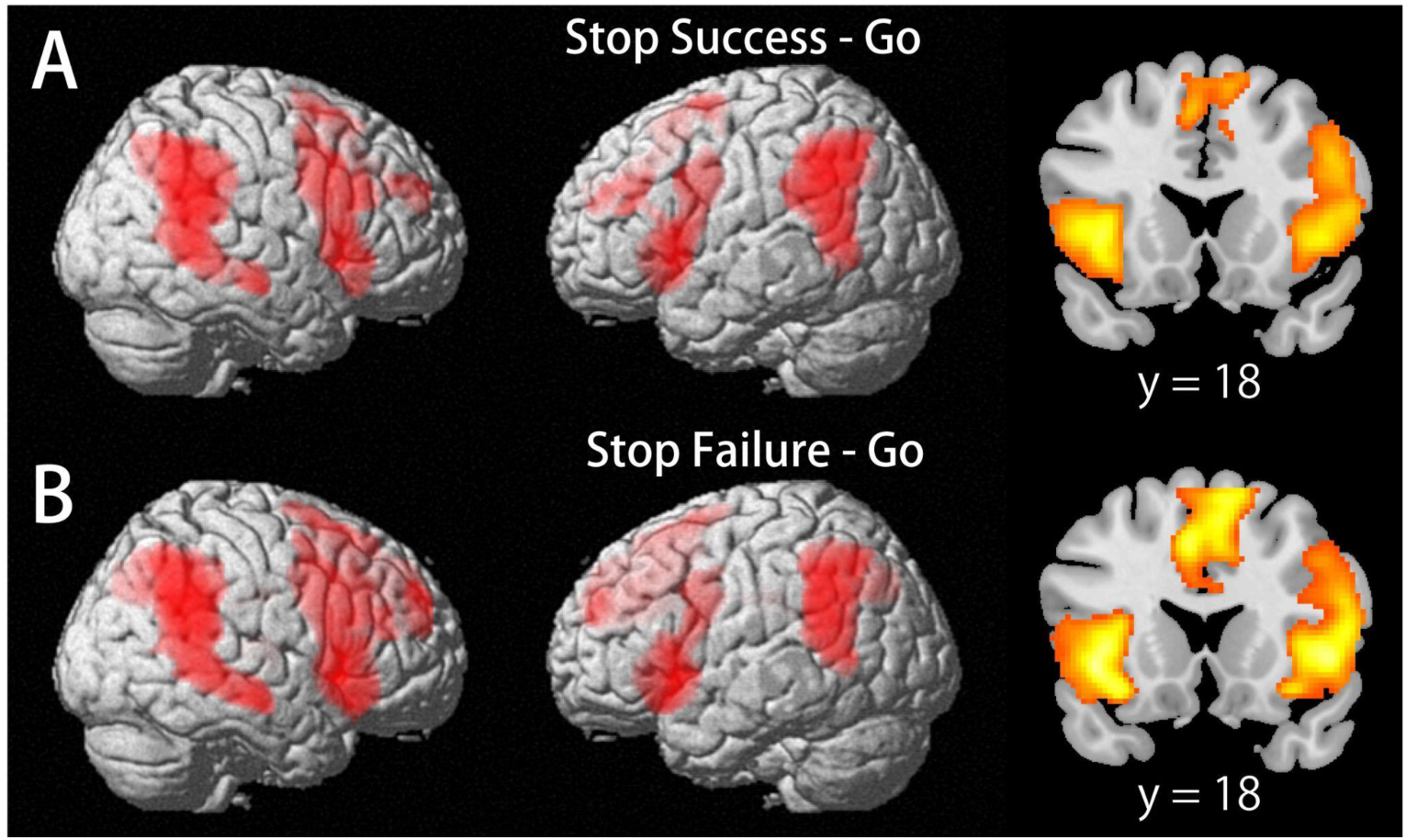
(A) Comparing Stop-success and Go trials, significant activity was noted in the bilateral parietal and frontal regions, insulae, and medial frontal gyri (detailed data presented in Table 4). (B) Analysing Stop-failure versus Go trials showed activity in the anterior cingulate cortex, in addition to the same regions as described above (Table 5).

**Table 4.** Brain activity in the contrast of Stop-success vs Go. All peak voxels are significant at p < 0.001, uncorrected with cluster-level correction (FWE p < 0.05) for the whole brain. *, FWE voxel-level corrected p < 0.01. **, This cluster extended bilaterally. FWE, family-wise error; Inf, Infinity.

| Cluster Size | Locations (Brodmann area) | x | y | z | Z-Value |
| --- | --- | --- | --- | --- | --- |
| 3629 | R. Superior Temporal Gyrus (22) | 64 | -48 | 18 | Inf* |
|  | R. Supramarginal Gyrus (40) | 60 | -54 | 30 | 7.80* |
|  | R. Inferior Parietal Lobule (40) | 64 | -42 | 26 | 6.94* |
| 2644 | L. Supramarginal Gyrus (40) | -62 | -50 | 32 | 7.76* |
|  | L. Supramarginal Gyrus (40) | -60 | -52 | 22 | 7.37* |
|  | L. Inferior Parietal Lobule (40) | -52 | -46 | 42 | 6.94* |
| 2917 | R. Inferior Frontal Gyrus (9) | 42 | 6 | 32 | 7.06* |
|  | R. Inferior Frontal Gyrus (47) | 42 | 20 | 0 | 5.72* |
|  | R. Inferior Frontal Gyrus (47) | 32 | 24 | -14 | 5.38* |
| 2445 | L. Inferior Frontal Gyrus (9) | -46 | 6 | 26 | 6.69* |
|  | L. Insula (13) | -38 | 20 | 0 | 6.47* |
|  | L. Inferior Frontal Gyrus (47) | -48 | 14 | 4 | 6.16* |
| 1175** | R. Superior Frontal Gyrus (6) | 12 | 10 | 66 | 5.67* |
|  | R. Superior Frontal Gyrus (6) | 8 | 22 | 56 | 4.99 |
|  | L. Medial Frontal Gyrus (8) | −4 | 16 | 50 | 4.77 |
| 240 | R. Middle Frontal Gyrus (10) | 24 | 52 | 26 | 5.12* |
|  | R. Middle Frontal Gyrus (10) | 30 | 42 | 30 | 3.69 |
| 256 | L. Superior Frontal Gyrus (6) | −16 | 2 | 66 | 4.48 |
|  | L. Middle Frontal Gyrus (6) | −28 | −2 | 54 | 3.30 |

**Table 5.**
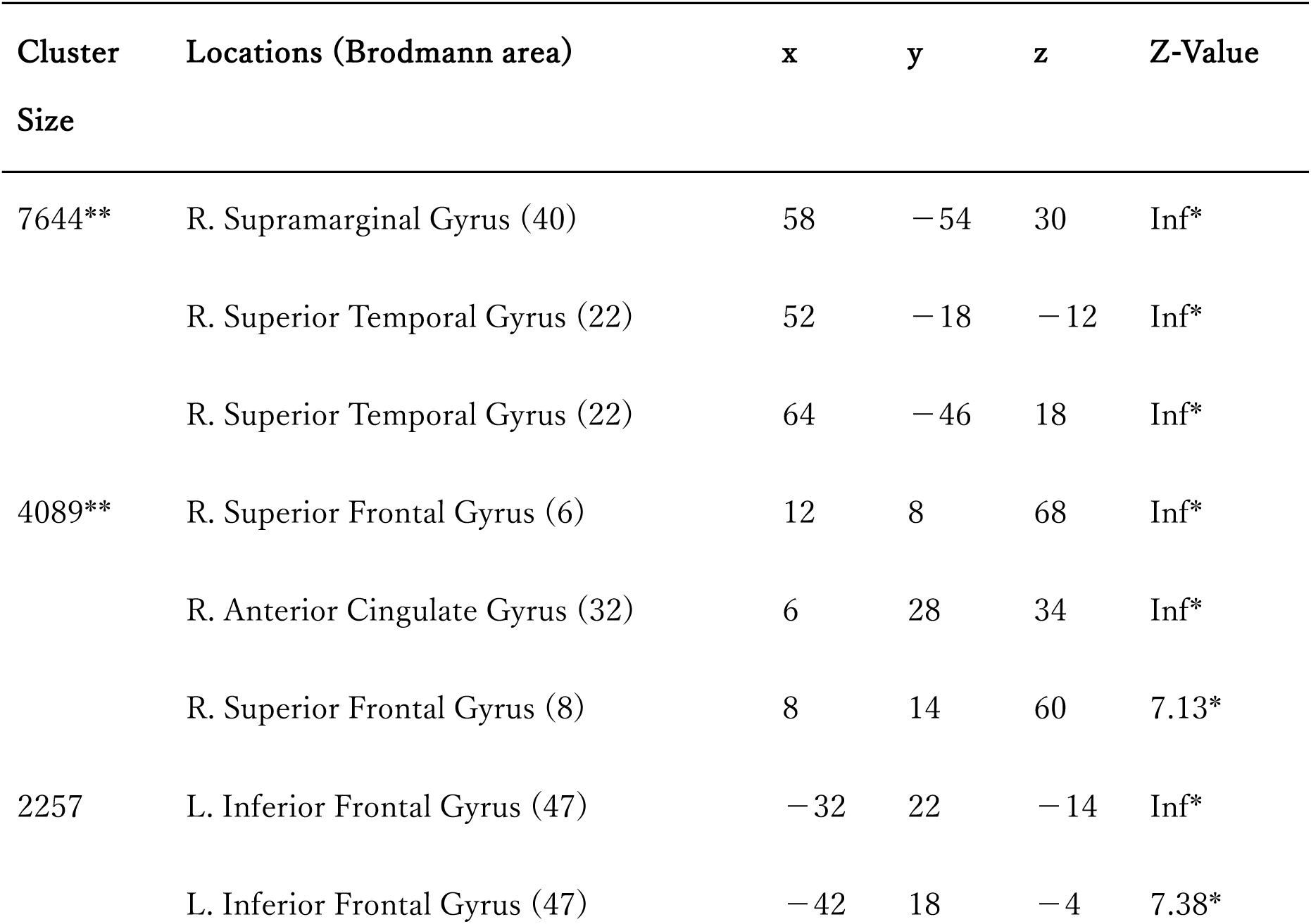
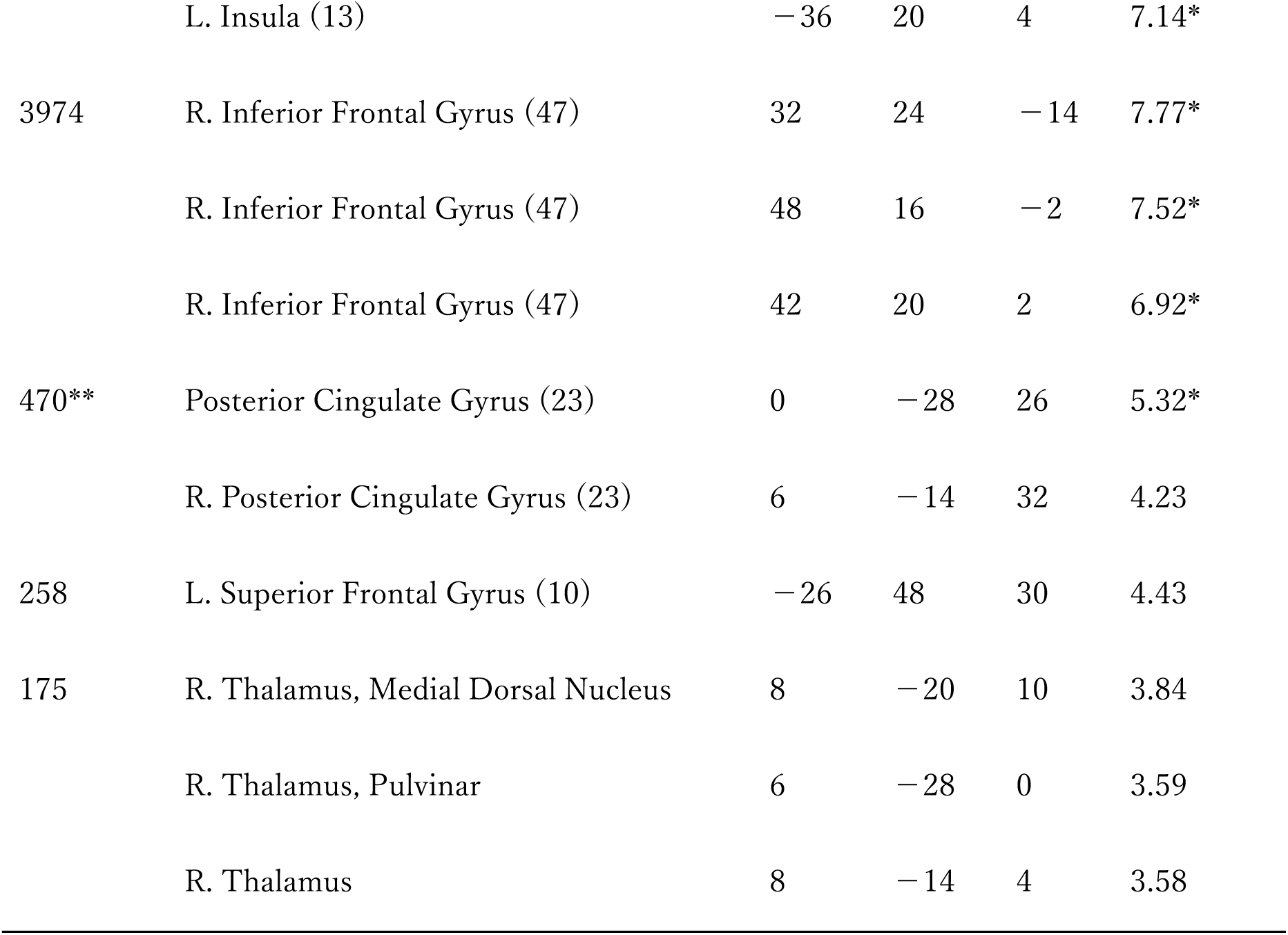
Brain activity in the contrast of Stop-failure vs Go. All peak voxels are significant at p < 0.001, uncorrected with cluster-level correction (FWE p < 0.05) for the whole brain. *, FWE voxel-level corrected p < 0.01. **, This cluster extended bilaterally. FWE, family-wise error; Inf, Infinity.

To investigate whether alcohol would enhance the difference in neural activity between SS and Go in some brain areas, an interaction analysis of (alcohol vs placebo) × (SS vs Go) was performed. However, no brain area was identified by this contrast in the whole brain, with a corrected threshold of p = 0.05. To test the specificity of the effect of alcohol on SF-related activity, an interaction analysis of (alcohol vs placebo) × (SF vs Go) was performed. The rIFC and the temporal pole were significantly activated only on the right side (Fig. 3A, Table 6). The percent signal change in this region showed SF activity specific to the alcohol condition (Fig. 3B). In addition, we investigated the brain regions where alcohol decreased the difference in neural activity between SS/SF and Go. Neither the combination of (placebo vs alcohol) × (SS vs Go) nor (placebo vs alcohol) × (SF vs Go) revealed any significant interaction in any area of the whole brain with a corrected threshold of p = 0.05.

**Figure 3:**
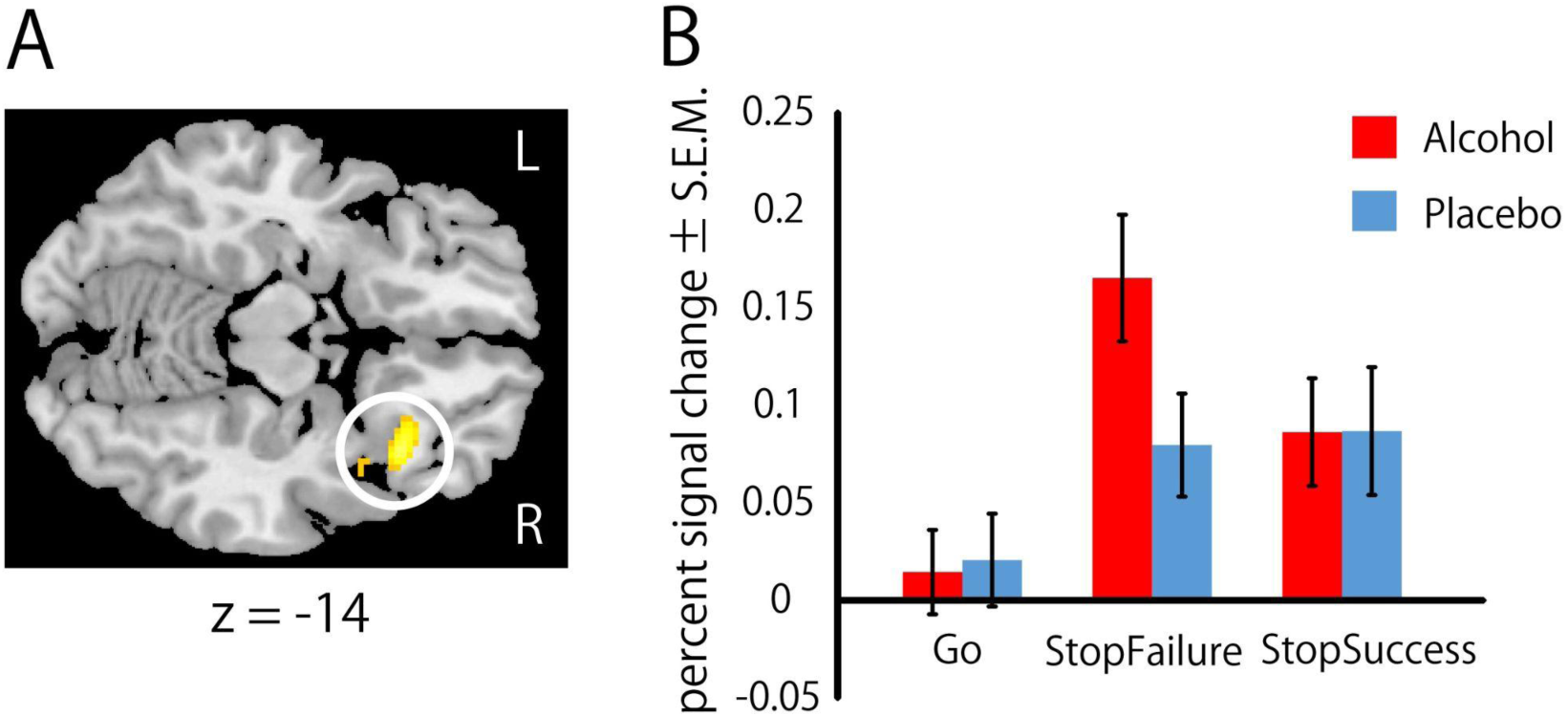
A) To examine how specifically alcohol impacts the activity related to Stop-failures (SFs), we conducted an interaction analysis using (alcohol vs placebo) × (SF vs Go). We found that the right inferior frontal cortex (rIFC) showed significant activation. B) The percent signal change in the rIFC in SF increases in the alcohol condition specifically. S.E.M., standard error of the mean.

**Table 6.** Brain activity in the interaction of (alcohol vs placebo) × (Stop-failure vs Go). All peak voxels are significant at p < 0.001, uncorrected with cluster-level correction (FWE p < 0.05) for whole brain. *, FWE voxel-level corrected p < 0.01. FWE, family-wise error.

| Cluster Size | Locations (Brodmann area) | x | y | z | Z-Value |
| --- | --- | --- | --- | --- | --- |
| 242 | R. Superior Temporal Gyrus (38) | 44 | 18 | -22 | 5.43* |
|  | R. Inferior Frontal Gyrus (47) | 36 | 24 | -14 | 4.39 |
|  | R. Inferior Frontal Gyrus (47) | 26 | 34 | -6 | 3.60 |

### Correlation between rIFC activity and PES

To investigate whether rIFC activity in the SF under alcohol conditions was related to PES under alcohol conditions, we conducted Pearson’s correlation analysis. There was no significant correlation between these variables (r = 0.177, t_15_ = 0.695, p = 0.498).

## DISCUSSIONS

In the present study, we investigated the effect of legally permissible low-level alcohol administration (0.15 mg/L BrAC) on behaviour including EMG and neural processing of response inhibition using SST. In the alcohol condition, Go RTs increased, and the prEMG frequency and normalised EMG magnitude (only in right arrow trials) of successful stops decreased. The increased Go RTs and decreased prEMG signals in the alcohol condition may reflect a delay in judgment (left or right arrow) and/or the participants’ careful manner of concentrating on the experiment. Since PES was significantly prolonged in the alcohol condition but not in the placebo condition, the time taken for behavioural adjustment after unsuccessful stopping (Li et al. 2006) may have increased due to alcohol consumption.

Go RTs were significantly prolonged and average SSD showed trend of increase in the alcohol condition compared with those in the placebo condition. Considering the almost constant SSRT in both conditions, this prolongation of Go RTs must be tightly linked to increased PES (17.4 ms vs. 10.6 ms) and average SSD (177.5 ms vs. 160.1 ms) under the alcohol condition. Previous reports have suggested that PES reflects behavioural adjustment after unsuccessful stopping (Li et al. 2006; Duann et al. 2009; Winkler et al. 2013; Manza et al. 2016; Zhang et al. 2017). Therefore, delayed Go RTs in the alcohol condition may also imply that behavioural adjustment took longer for Go trials after the SF trials. However, in a previous study, participants with relatively high alcohol intoxication did not show a significant increase in Go RTs during the SST (Gan et al. 2014). The low alcohol intoxication in the present study may have drawn proactive attention and/or urged caution caused by the awareness of drinking, while the high alcohol intoxication may not have raised attention to the SST.

Regarding the SSRT, there was no significant effect of alcohol consumption in the present study. A previous study reported that SSRT increased under a high dose of alcohol but did not increase under a low dose of alcohol (de Wit et al. 2000). Our study replicated part of this previous study; that is, a low dose of alcohol did not affect SSRT. The SST is a race between the Go and Stop units (Logan and Cowan 1984; Verbruggen and Logan 2008; Schall et al. 2017). Therefore, the Stop unit may not have been affected by a low alcohol dose.

rIFC activity increased under alcohol conditions during SF trials, as shown in Fig. 3. To infer the role of the rIFC in this study, we examined two possible interpretations of this observation for consistency with previous data; either the increased activity of the rIFC led to the failure of stopping, or the failure of stopping resulted in increased rIFC activity. First, let us assume that increased activity of the rIFC leads to failure to stop. Failure to stop implies that the activation related to the Stop unit is weaker than that related to the Go unit. Therefore, an increase in rIFC activity was associated with a decrease in Stop unit activation. However, this is inconsistent with previous studies showing that increased rIFC activity was associated with increased activation of the Stop unit (for review see Aron et al. 2014).

Next, we hypothesised that increased activity in the rIFC results from SF. In this study, significant PES after SF occurred only under alcohol conditions. Previous reports have suggested that the PES reflects behavioural adaptation following failure to stop (Li et al. 2006; Duann et al. 2009; Winkler et al. 2013; Manza et al. 2016; Zhang et al. 2017). Additionally, studies using repetitive TMS indicated that the rIFC during the SST may be involved in reactive control (Zandbelt et al. 2013). These results are consistent with the second interpretation.

The normalised EMG magnitude during failed Stop trials was less than one. This measure was based on the EMG magnitude during the Go trials, which was set to one. Because we defined the normalised EMG magnitude during failed Stop trials as the SF EMG power divided by the Go EMG power, we would expect the normalised EMG magnitude during failed Stop trials to be one if the Stop signal did not affect muscle contraction. Considering these points, muscle contraction during failed Stop trials is suggested to be influenced by Stop signals.

In this study, the correlation coefficient between rIFC activity and PES was positive, although not significant. This finding might support the hypothesis that the rIFC is involved in behavioural adaptation following the failure to stop. Further research with a larger sample size is necessary to robustly demonstrate this hypothesis.

The task design used in this study was the same as that used in a previous study (Tabu et al. 2011). Tabu et al. reported significant activity in the bilateral IFC/insula, right dorsolateral prefrontal cortex, pre-supplementary motor area, and parietal cortex when comparing SS and Go trials. In our study, significant activity in these same regions was observed in the SS vs Go contrast, as shown in Table 4 and Fig. 2. This activity during response inhibition has also been observed in other studies (Aron and Poldrack 2006; Cai and Leung 2009; Chikazoe et al. 2009; Duann et al. 2009; Sharp et al. 2010). Thus, our study replicated these findings. Moreover, regarding behaviour and EMG, a previous meta-analysis (Raud et al. 2022) indicated an inverse relationship between Go RTs and prEMG frequency, which was also replicated in our study.

A limitation of this study is that some participants were able to discern whether the beverage contained alcohol based on stimuli such as the smell of alcohol. This may have led to the intentional adjustment of the speed-accuracy trade-off. We believe that the activity in the rIFC observed in this study was involved in adjusting the speed-accuracy trade-off following failed stop attempts. However, the extent to which this adjustment is intentional and the degree to which intentionality is associated with rIFC activity remain unclear. Future research should explicitly separate these intentions from the task design.

In conclusion, our results illustrate that even a low concentration of alcohol (0.15 mg/L) changes human behaviour and brain activity. Because a BrAC of 0.15 mg/L is the lower limit of the regulation for driving in Japan, it is important to review the regulation.

## ACKNOWLEDGEMENTS

We would like to thank Editage (www.editage.jp) for English language editing.

## REFFERENCES

Aron AR, Fletcher PC, Bullmore ET, Sahakian BJ, Robbins TW. Stop-signal inhibition disrupted by damage to right inferior frontal gyrus in humans. Nat Neurosci. 2003:6(2):115‒116.

Aron AR, Poldrack RA. Cortical and subcortical contributions to stop signal response inhibition: role of the subthalamic nucleus. J Neurosci. 2006:26(9):2424‒2433.

Aron AR, Robbins TW, Poldrack RA. Inhibition and the right inferior frontal cortex: one decade on. Trends Cogn Sci. 2014:18(4):177‒185.

Badry R, Mima T, Aso T, Nakatsuka M, Abe M, Fathi D, Foly N, Nagiub H, Nagamine T, Fukuyama H. Suppression of human cortico-motoneuronal excitability during the stop-signal task. Clin Neurophysiol. 2009:120(9):1717‒1723.

Cai W, Leung HC. Cortical activity during manual response inhibition guided by color and orientation cues. Brain Res. 2009:1261(March):20‒28.

Chambers CD, Bellgrove MA, Gould IC, English T, Garavan H, McNaught E, Kamke M, Mattingley JB. Dissociable mechanisms of cognitive control in prefrontal and premotor cortex. J Neurophysiol. 2007:98(6):3638‒3647.

Chambers CD, Bellgrove MA, Stokes MG, Henderson TR, Garavan H, Robertson IH, Morris AP, Mattingley JB. Executive ʻbrake failure’ following deactivation of human frontal lobe. J Cogn Neurosci. 2006:18(3):444‒455.

Clark L, Blackwell AD, Aron AR, Turner DC, Dowson J, Robbins TW, Sahakian BJ. Association between response inhibition and working memory in adult ADHD: a link to right frontal cortex pathology? Biol Psychiatry. 2007:61(12):1395‒1401.

Duann JR, Ide JS, Luo X, Ray Li CS. Functional connectivity delineates distinct roles of the inferior frontal cortex and presupplementary motor area in stop signal inhibition.” J Neurosci. 2009:29(32):10171‒ 10179.

Evans AC, Collins DL, Mills SR, Brown ED, Kelly RL, Peters TM. 3D Statistical Neuroanatomical Models from 305 MRI Volumes. In 1993 IEEE Conference Record Nuclear Science Symposium and Medical Imaging Conference, 1993:1813‒17. IEEE.

Friston KJ, Holmes AP, Worsley KJ, Poline JP, Frith CD, Frackowiak RSJ. Statistical parametric maps in functional imaging: a general linear approach. Hum Brain Mapp. 1994:2(4):189‒210.

Gan G, Guevara A, Marxen M, Neumann M, Jünger E, Kobiella A, Mennigen E, Pilhatsch M, Schwarz D, Zimmermann US, Smolka MN, Alcohol-induced impairment of inhibitory control is linked to attenuated brain responses in right fronto-temporal cortex. Biol Psychiatry. 2014:76(9):698‒707.

Hajcak G, McDonald N, Simons RF. To err is autonomic: error-related brain potentials, ans activity, and post-error compensatory behavior. Psychophysiology. 2003:40(6):895‒903.

Koelega HS. Alcohol and vigilance performance: a review. Psychopharmacology (Berl). 1995:118(3):233‒ 249.

Leung HC, Cai W. Common and differential ventrolateral prefrontal activity during inhibition of hand and eye movements. J Neurosci. 2007:27(37):9893‒9900.

Levy I, Lazzaro SC, Rutledge RB, Glimcher PW. Choice from non-choice: predicting consumer preferences from blood oxygenation level-dependent signals obtained during passive viewing. J Neurosci. 2011:31(1):118‒125.

Li CS, Huang C, Constable RT, Sinha R. Imaging response inhibition in a stop-signal task: neural correlates independent of signal monitoring and post-response processing. J Neurosci. 2006:26(1):186‒192.

Loeber S, Duka T. Acute alcohol impairs conditioning of a behavioural reward-seeking response and inhibitory control processes--implications for addictive disorders. Addiction. 2009:104(12):2013‒ 2022.

Logan GD, Cowan WB, Davis KA. On the ability to inhibit simple and choice reaction time responses: a model and a method. J Exp Psychol Hum Percept Perform. 1984:10(2):276‒291.

Logan GD, Cowan WB. On the ability to inhibit thought and action: a theory of an act of control. Psychol Rev. 1984:91(3):295‒327.

Manza P, Hu S, Chao HH, Zhang S, Leung HC, Li CR. A dual but asymmetric role of the dorsal anterior cingulate cortex in response inhibition and switching from a non-salient to salient action. Neuroimage. 2016:134(July):466‒474.

Marinkovic K, Halgren E, Maltzman I. Arousal-related P3a to novel auditory stimuli is abolished by a moderately low alcohol dose. Alcohol Alcohol. 2001:36(6):529‒539.

Mulvihill LE, Skilling TA, Vogel-Sprott M. Alcohol and the ability to inhibit behavior in men and women. J Stud Alcohol. 1997:58(6):600‒605.

Rabbitt PM. Errors and error correction in choice-response tasks. J Exp Psychol. 1966:71(2):264‒272.

Raud, L, Thunberg C, Huster RJ. Partial response electromyography as a marker of action stopping. eLife. 2022:11:e70332

Rubia K, Smith AB, Brammer MJ, Taylor E. Right inferior prefrontal cortex mediates response inhibition while mesial prefrontal cortex is responsible for error detection. Neuroimage. 2003:20(1):351‒358.

Scheel JF, Schielke K, Lautenbacher S, Aust S, Kremer S, Wolstein J. Low-dose alcohol effects on attention in adolescents. Zeitschrift Für Neuropsychol. 2013:24(2):103‒111.

Tabu H, Mima T, Aso T, Takahashi R, Fukuyama H. Functional relevance of pre-supplementary motor areas for the choice to stop during stop signal task. Neurosci Res. 2011:70(3):277‒284.

Tzambazis K, and Stough C. Alcohol impairs speed of information processing and simple and choice reaction time and differentially impairs higher-order cognitive abilities. Alcohol Alcohol. 2000:35(2):197‒201.

Verbruggen F, Aron AR, Stevens MA, Chambers CD. Theta burst stimulation dissociates attention and action updating in human inferior frontal cortex. Proc Natl Acad Sci U S A. 2010:107(31):13966‒ 13971.

Verbruggen F, Logan GD. Response inhibition in the stop-signal paradigm. Trends Cogn Sci. 2008:12(11):418‒424.

Winkler AD, Hu S, Li CS. The influence of risky and conservative mental sets on cerebral activations of cognitive control. International Journal of Psychophysiology. 2013:87(3):254261.

de Wit H, Crean J, Richards JB. Effects of d-amphetamine and ethanol on a measure of behavioral inhibition in humans. Behav Neurosci. 2000:114(4):830‒837.

Zandbelt BB, Bloemendaal M, Hoogendam JM, Kahn RS, Vink M. Transcranial magnetic stimulation and functional MRI reveal cortical and subcortical interactions during stop-signal response inhibition. J Cogn Neurosci. 2013:25(2):157‒174.

Zhang Y, Ide JS, Zhang S, Hu S, Valchev NS, Tang X, Li CS. Distinct neural processes support post-success and post-error slowing in the stop signal task. Neuroscience 2017:357(August):273‒284.

