## Supplementary material for "Low blood concentration of alcohol enhances activity related to stopping failure in the right inferior frontal cortex": Fig. S1

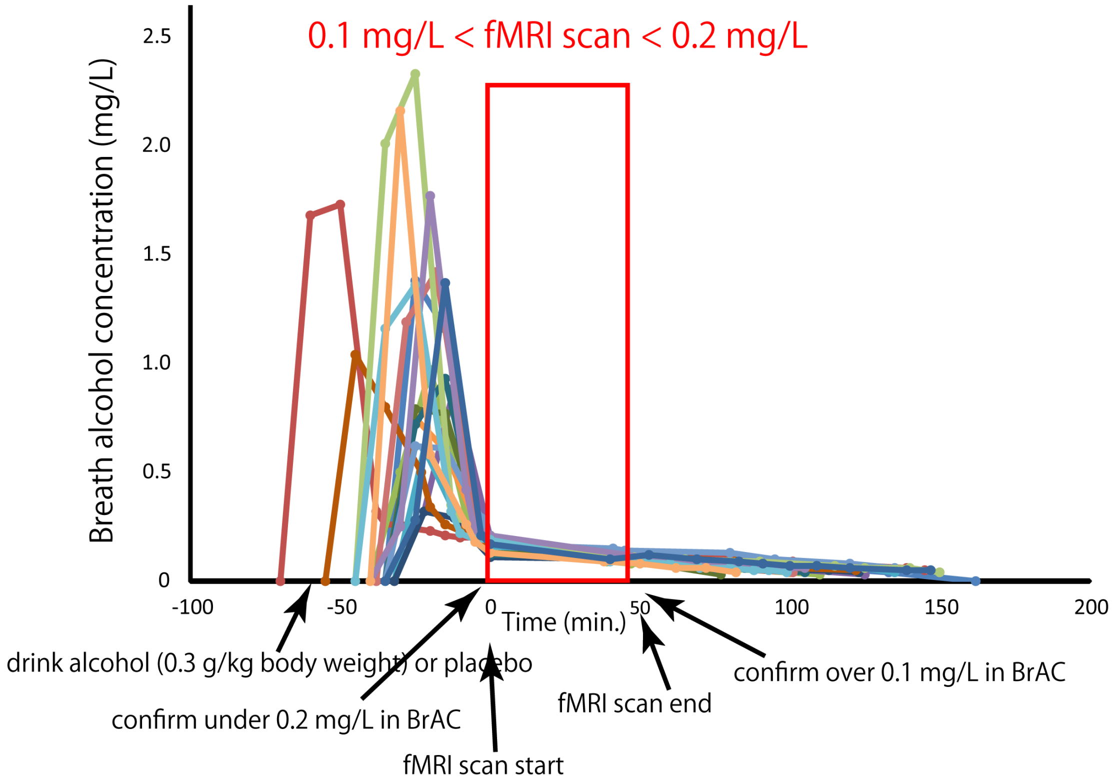


Fig S1

Time course of breath alcohol concentration (BrAC) for all participants (n = 17). We checked BrAC every five minutes and started the fMRI scan when BrAC dropped to 0.20 mg/L or lower. To be included in the study, participants had to have a post-scan BrAC of 0.10 mg/L or greater.
